# Muscle synergies to reduce the number of electromyography channels in neuromusculoskeletal modelling: a pilot study

**DOI:** 10.1101/2024.09.24.614668

**Authors:** M. Romanato, F. Spolaor, Z. Sawacha

**Affiliations:** Department of Information Engineering, University of Padova, Padova, Italy

**Keywords:** muscle synergies, neuromusculoskeletal modeling, rehabilitation engineering, motor control, translational research

## Abstract

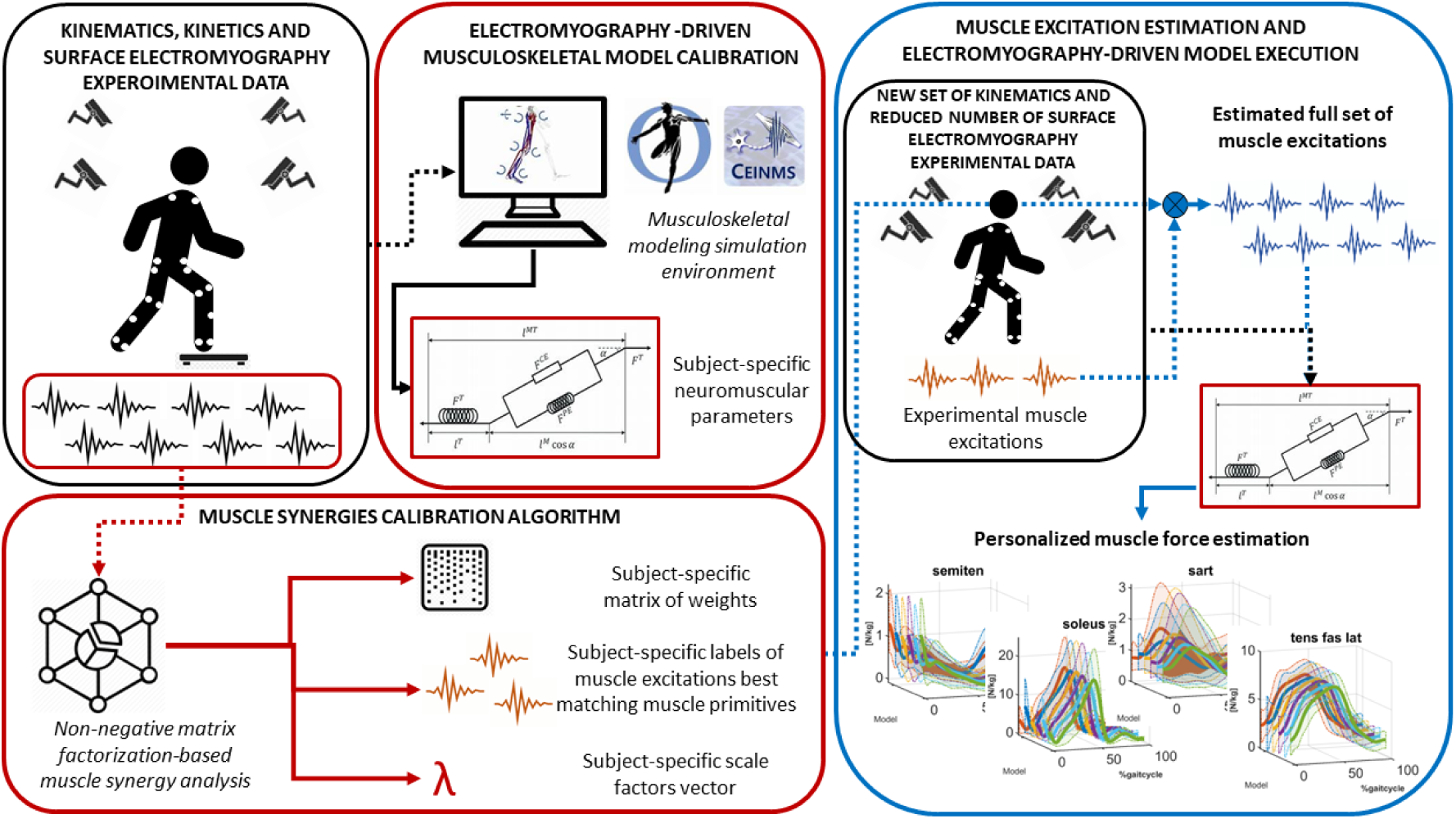

**Graphical abstract.** In the black squares, we present recorded experimental data. The black box on the left contains information about the experimental data used for calibration purposes, including kinematics, kinetics, and surface electromyography (sEMG) data. The black box on the right contains information about the experimental data used for testing the algorithm, which includes kinematics and a reduced set of sEMG data. In the red boxes, we depict two parallel calibration processes. The bottom red box outlines the muscle synergies calibration algorithm, which takes as input the sEMG recordings from the calibration set (black box on the left) and provides subject-specific parameters for reconstructing missing muscle excitations from a new set of collected sEMGs. The top red box illustrates the sEMG-driven musculoskeletal model calibration, which takes input from kinematics, kinetics, and sEMG data, yielding a calibrated model with subject-specific neuromuscular parameters. The blue box describes how subject-specific muscle forces are estimated from a new dataset with limited sEMG recordings. A complete set of muscle excitations is estimated based on these few recordings and the subject-specific parameters obtained in the muscle synergies calibration algorithm box. These excitations, along with the kinematics data, drive the calibrated musculoskeletal model obtained in the sEMG-driven musculoskeletal model calibration box. Dotted arrows represent inputs, and solid lined arrows represent outputs.

## 1. INTRODUCTION

Rehabilitation therapy has a significant importance in many motor disorders, with the purpose of maximizing functional ability and minimizing secondary complications through movement restoration [1]. Thus, there is a pressing need to find effective methods to improve quality-of-life in people with motor disorders. However, the long-term retention of its potential benefits remains to be formally determined due to the lack of objective methodologies (Peppe et al., 2007). Techniques to measure muscle strength regaining after physical intervention are mainly based on manual tests [3]. These are delivered in isometric and quasistatic conditions by a single trained operator, and thus might be either affected by unvoluntary biases, or by the lack of functional sensorimotor recovery characterization during dynamic tasks. Instrumented gait analysis is widely recognized to describe human motion in pathological conditions [4,5]. However, current motion capture techniques do not provide insights on the interplay between neural to muscles activation, nor on disrupted musculoskeletal functions *in vivo*.

Surface electromyography (sEMG)-driven models are powerful tools for personalized rehabilitation protocols. These approaches provide information on how neural command interplays with the mechanical output, through the definition of quantities such as muscle forces and activations in a simulation environment [6]. These models, informed via the experimentally measured subject’s muscle excitations, can be calibrated to provide a higher degree of personalization, and allows to capture the subject-specific neural-to-mechanical interplay. Through the estimation of quantitative and repeatable metrics, new sets of plausible biomechanical biomarkers can be defined, namely muscle forces, which can objectively report on the motor capacity of the patient, reducing the possible sources of variability affecting the clinical scales [7,8]. Despite the informativeness of those approaches, their use outside of laboratory settings remains hampered due to the high number of required sEMG signals [9]. Indeed, neuromusculoskeletal models are rarely adopted in clinical evaluations because of their requirement of expensive instruments (*i.e.*, stereophotogrammetry, force plates, sEMG) which can’t be applied in out-of-the-laboratory settings. A first attempt to propose a sEMG-driven neuromusculoskeletal model suitable for clinical use through a reduction of the needed input was recently made by the authors (Romanato et al., 2023). However, limitations of that work were acknowledged: *1)* the reduced input considered in that work consisted only of the combination of one set of agonist\antagonist muscles (*i.e.*, biceps femoris, rectus femoris, gastrocnemius lateralis, tibialis anterior), selected as the most assessed in clinical gait analysis [10,11]; *2)* the subject-specificity of motor control pattern captured via muscle synergy analysis was not considered. Muscle synergy analysis has been proposed to predict unmeasured muscle excitations from the measured signals [12–14]. They are defined by a time-varying primitive, and a time-invariant matrix of weights that describe how the primitives contribute to the excitation of all the considered muscles [15,16]. This mathematical tool could be adopted to reduce the number of input signals needed to drive neuromusculoskeletal models [17], aiding the translation of these methodologies from gait laboratories to clinical applications, or even to domestic environments.

The aim of the present study is to explore the reliability of a calibrated muscle-synergies approach to identify subject-specific motor control parameters to be employed in the prediction of unmeasured muscles excitations while considering few sEMG recordings as muscle primitives. Then, the new set of synthetized muscle excitations would be employed to drive a neuromusculoskeletal model to predict subject-specific joint torques and muscle forces of the lower-limb muscle groups. It was hypothesized that a minimal set of sEMG signals determined through muscle synergy analysis can be used to drive a neuromusculoskeletal model. Given the reduction of the modeling inputs, a validation of the proposed framework could be one of the first steps to provide personalized home-based rehabilitation treatments that will enable clinicians to perform online tracking of both disease progression and treatment outcomes.

## 2. METHODS

### 2.1. Participants

Kinematics, kinetics and sEMG data of 4 non-disabled adults (female\male = 1\3; age = 60.0 ± 2.1 years; body mass index = 26.7 ± 4.1 kg/m^2^) have been recorded while walking at the BioMovLAB (Department of Information Engineering, University of Padova, Padova, Italy). Participants’ age was chosen to potentially match the one in which commonly neurodegenerative disorders onset [18]. Participants were asked to perform several walking trials barefoot at their preferred walking speed. To be included in the present study, participants were able to walk independently, did not have pathologies that prevent safe mobility, and were free of musculoskeletal, neurologic, cardiopulmonary, and other systemic disorders. They were enrolled to the study after comprehending and signing written informed consent to the experimental protocol (Ethic Committee, number 1001 P -21/11/2005, University Policlinic of Padova [19]).

### 2.2. Experimental setup

A 6-camera stereophotogrammetric system (60 Hz, SMART-E, BTS Bioengineering, Italy) was used to capture the three-dimensional markers’ trajectory, synchronized with two inground force plates (960 Hz, fp4060, Bertec Corporation, USA) recording ground reaction forces and a 16-channels sEMG system (1000 Hz, BTS-Free1000, BTS Bioengineering, Italy) recording muscle’s electrical activity. Participants were instrumented with retroreflective spherical markers, placed according to the IORgait marker set [20], and with 15 sEMG probes, positioned on the muscle belly of 15 ipsilateral lower-dominant-limb muscles following the SEMIAM guidelines [21] after appropriately shaving and cleaning the skin. Electrodes (Ø=24mm, Hydrogel/Ag/AgCl, Kendall Arbo H124SG, Ireland) were located 1 cm apart. The electrical activity of the gluteus medius, gluteus maximus, adductor longus, tensor fasciae latae, sartorius, bicep femoris, semitendinosus, rectus femoris, vastus medialis, vastus lateralis, gastrocnemius lateralis, gastrocnemius medialis, soleus, peroneus longus and tibialis anterior was collected.

### 2.3. Data processing

Twelve complete right walking cycles per participant were considered. The trials were included in the analysis if the right foot naturally landed in one force plate, and they were segmented when identified by two consecutive heel strikes [22]. MOtoNMS [23] was used to filter and process markers trajectories and ground reaction forces to obtain an OpenSim-compatible format. OpenSim (v. 4.4, SimTK) was employed to anisotropically scale a generic musculoskeletal model (gait2349, [24]) that best match the subject’s anthropometrics and inertial properties, considering subject’s marker trajectories during static pose. The batch OpenSim processing scripts (BOPs 0.9, SimTK [25]) were adopted to run the inverse kinematics, inverse dynamics, and muscle analysis tools to compute joint angles, joint moments and muscle-tendon units kinematics respectively. MOtoNMS was also adopted to process the sEMG signals and obtain their linear envelope after band pass filtering (30-300 Hz, 4^th^ order Butterworth), full wave rectification and low pass filtering (6 Hz, 4^th^ order Butterworth). The signals were normalized to the maximum sEMG value recorded across all trials.

Per each participant, the dataset was randomly split in two: six trials were considered for calibration, while the remaining six trials were considered for validation, accordingly with previously done in similar studies [12]. Custom made MATLAB (v. R2021b, Mathworks) scripts were used for data processing.

### 2.4. Muscle excitation prediction via synergy-based parameter calibration

This process involved two main steps: a training step (*synergy-based parameter calibration*), where the set of representative muscles and subject-specific parameters were defined, and a testing step where excitation from unmeasured muscles were predicted and compared with the experimental excitations (*unmeasured excitation prediction*). In Figure 1 is reported a schematic representation of the process.

**Figure 1.**
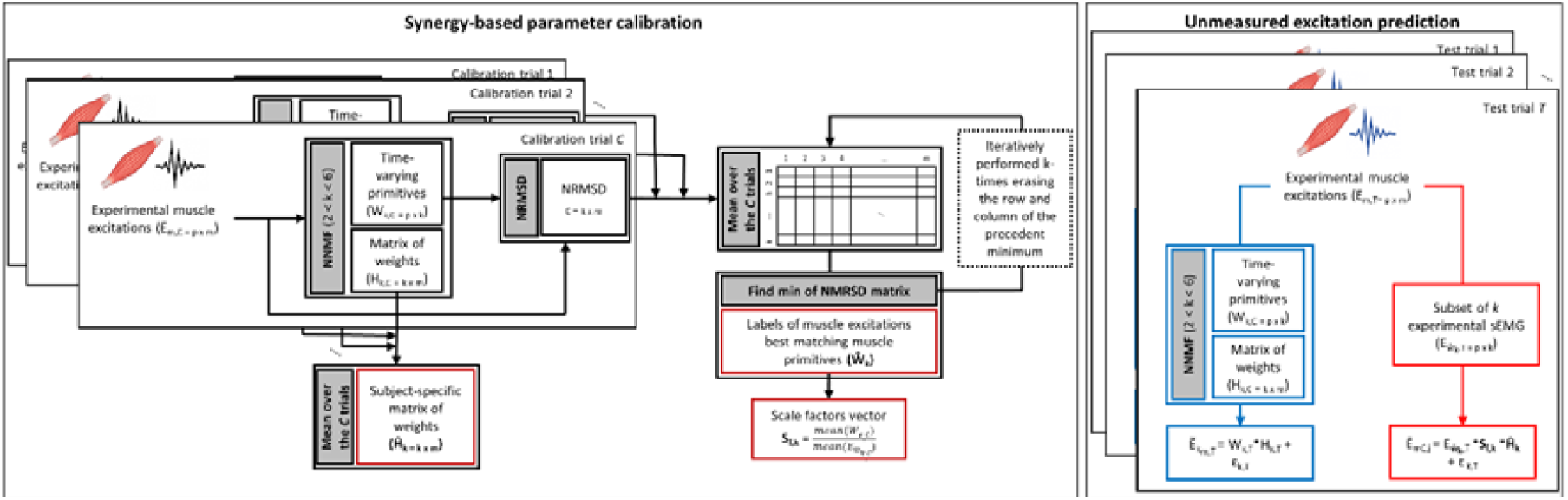
Schematic representation of the Synergy-based parameter calibration on the left. Schematic representation of the unmeasured excitation prediction process (red arrows) on the right. Blue arrows indicated the process used for comparison.

#### Synergy-based parameter calibration

The calibration set was used to extract the muscle synergies primitives (*W_k,C_*) and weights matrix (*H_k,C_*) through non-negative matrix factorization (NNMF) methods, where *k* was the factorization number. This number varied between 2 and 6, as typically reported in literature [16,26], and corresponded to the number of muscle synergies considered for the extraction from the whole set of 15 sEMG signals *W_k,C_* and *H_k,C_* were assumed to be subject and trial specific, so that *C=1, …, n*, with *n* total number of the calibration trials. The objective function minimized by the NNMF method was:

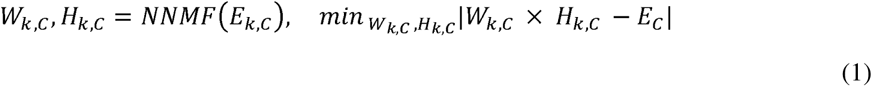

Where *E_m,C_* is the matrix of the *m*=15 sEMG signals of the *C*^th^ trial time-normalized over *p*=100 samples, thus *E_m,C_* has *p×m* dimensions.*E_m,C_* has *p×k* dimensions, while *W_k,C_* has *k×m* dimensions. The normalized root mean squared difference (*NRMSD*) between each experimental sEMG signals *m* and each *k^th^* synergy vector from *W_k,C_* was computed as follows,

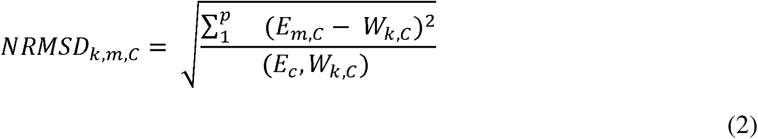

*NRMSD_k,,m,C_* was a 3-dimensional matrix containing all the *NRMSD* values for each calibration trial. The *NRMSD* average along the third dimension was used as metric to determine which *k*-subset of the *m* experimental sEMG signals better resembled, 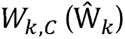: firstly, the lowest value of *NRMSD* determined the *m*-*k* pair with the highest level of agreement, then the corresponding column and row of the *NRMSD* matrix were deleted from the research grid, and the second lowest value of *NRMSD* identified to determine the second *m-k* pair. The process was repeated iteratively *k* times. Finally, 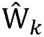 contains the subject-specific labels of the sEMG signals with the highest similarity with respect to the retrieved motor primitives. To account for the differences between the two signals a vector of scaling factors *S_f,k_* was defined as the ratio between the average across each calibration trial of *W_k,C_* and the average across each calibration trial of the 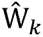 signals extracted from *E_m,C_*, defined as 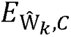.

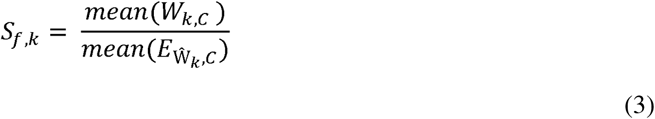

*S_f,_* is a subject-specific vector containing *m* scale factors, one for each sEMG signal. Finally, the average of the *H_k,C_* across each trial was considered as subject-specific weights matrix 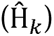. Therefore, once calibration was performed, the terms allowing personalization were: the subject-specific muscle tags 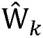 the vector *S_f,k_* and the matrix 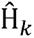.

#### Unmeasured excitation prediction

In the testing step, *E_m,T_* was the matrix of the *m*=15 sEMG signals of *T*^th^ trial, being *T=1, …, i,* with *i* total number of the testing trials. 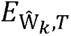, was the matrix of the *T*^th^ trial that considered a subset of the *k=2, …, 6* experimental sEMG signals defined by 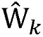, and therefore has dimensions of p×k. The *k–m* sEMG signals that were not considered by were treated as “unmeasured”. Thus, to reconstruct the original matrix *E_m,T_* the following formula was applied:

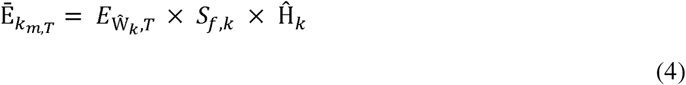

where Ē*_m,T_* was the reconstruction of *E_m,T_* obtained using a subset of *k* experimentally recorded sEMG signals.

In the same way, the actual trial-specific primitives (*W_k,T_*) and vector of weights (*H_k,T_*) have been extracted via NNMF from *E_m,T_* as in Equation 1, for *k=2, …, 6*. Therefore, they were used to reconstruct *E_m,T_* applying

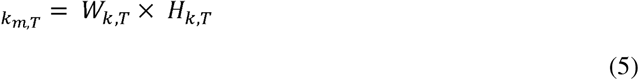

where *k_m,T_* was the reconstruction of *E_m,T_* obtained using *k* muscle synergies.

CEINMS [27] was used as modeling environment for the modeling of neuromusculoskeletal system. This consisted of two processes, (*model calibration* and *model execution*), that were performed in parallel with the ones described in Section 2.4 respectively. Each model consisted of 24 muscle tendon units (MTUs) spanning over four degrees of freedom (*i.e.*, hip flexion-extension, hip ab-adduction, knee flexion-extension, ankle dorsi-plantarflexion).

#### Neuromusculoskeletal model calibration

Six calibration trials were used in this process to refine within physiological bounds the neuromuscular parameters of the model through minimization of the error between experimental and model predicted joint moments [27,28]: tendon slack length [0.85, 1.15], optimal fiber length [0.85, 1.15], non-linear shape factor [-2.999, -0.001], strength coefficient [0.5, 3] and activation non-linearity constants [-0.95, -0.05]. MTUs kinematics, joint moments and experimental sEMG signals were used as input for the process. Experimental excitations were mapped over the 24 MTUs as in Table 1.

**Table 1.**
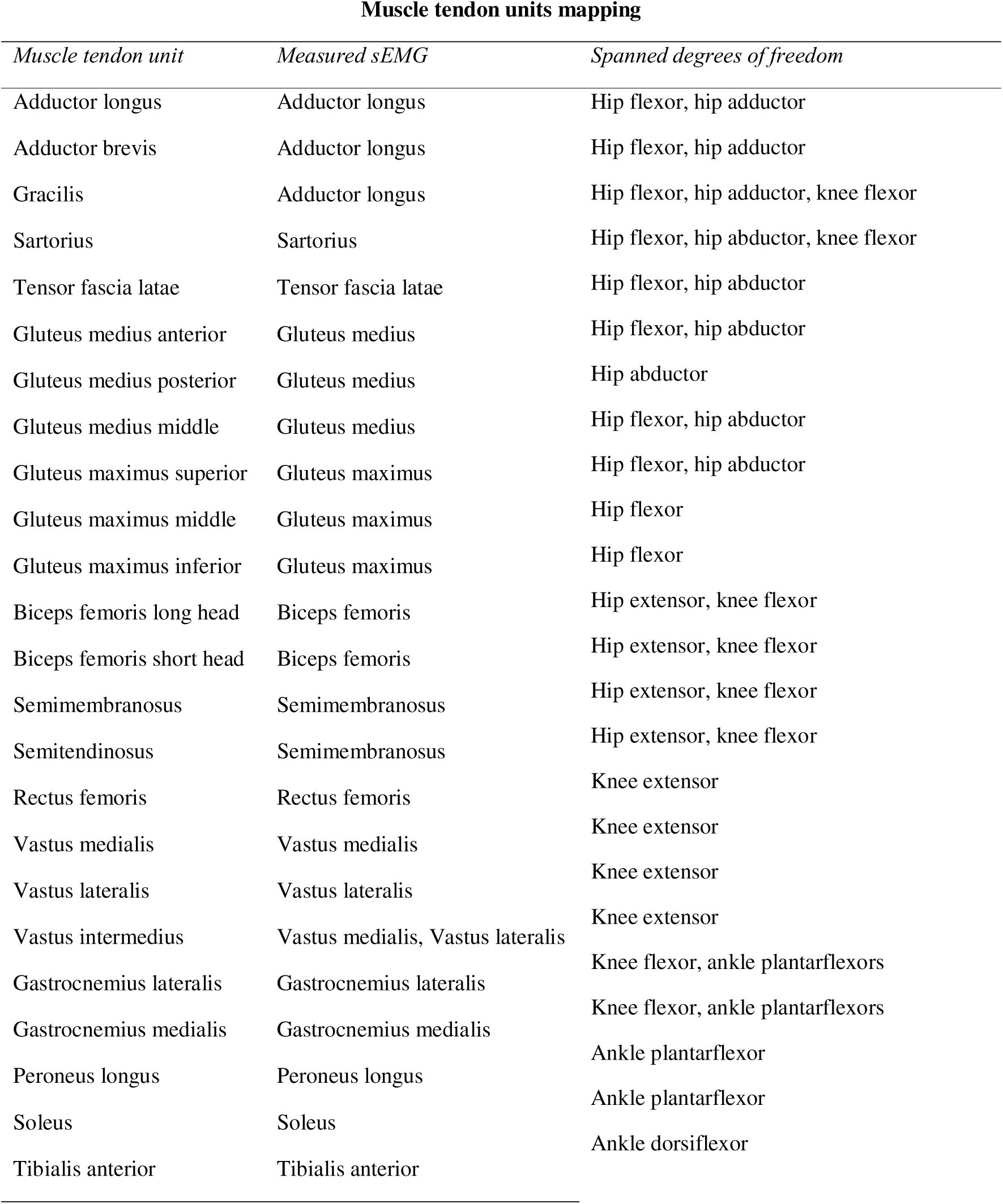
MTUs considered in the neuromusculoskeletal models, the recorded sEMG used to map their activation dynamics and the degree of freedom spanned by the MTU.

#### Neuromusculoskeletal model execution

Six testing trials per subject were used to estimate joint moments and muscle forces using an open-loop sEMG-driven approach. To investigate how the excitation reconstruction method proposed in Section 2.4 affects the outcomes of the model execution, Ē*_m,T_* were used as input to feed the model, thus obtaining six different outcomes per each different trial (five corresponding to the different realizations of Ē*_m,T_*, being *k*=2, …, 6, and one corresponding to experimental sEMG signal *E_m,T_*).

### 2.5. Data Analysis

Root mean squared difference (RMSD), determination coefficient (R^2^) variance accounted for (VAF) between the experimental sEMG signals (*E_m,T_*) and the simulated (Ē*_m,T_*, or *k_m,T_*,) excitations were employed to quantify the matching of magnitude, shape, and reconstruction accuracy respectively of each estimated curve. NRMSD, R^2^ and mean absolute error (MAE) were calculated between the experimental joint moments from inverse dynamics and the model predicted ones. RMSD, R^2^ were also adopted to account for differences in magnitude and shape of the estimated muscle forces, considering as *gold standard* the model driven by the experimental sEMG signals (*E_m,T_*,). R^2^ was considered weak if R^2^≤0.35, moderate if 0.35<R^2^≤0.67, strong if 0.67<R^2^≤0.9, very strong if 0.9≤R^2^ [29].

ANOVA with a Tukey post-hoc analysis on MAE, NMRSD and R^2^ for the joint torques and muscle forces was performed to assess the influence of *k* in reconstructing the full set of experimental recordings, setting the significance level at *p*<0.05 (v. 4.1.0, RStudio).

## 3. RESULTS

### 3.1. Experimental sEMG reconstruction accuracy

Subject-specific muscle excitations (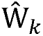) defined in the calibration part that has been more frequently used to reproduce the unmeasured muscle excitations have been summarized in Figure 2. Across the *k*-conditions, sEMG signals that have been recognized to better reproduce motor control patterns are tibialis anterior, soleus, peroneus longus, gastrocnemius medialis, semitendinosus and the adductor longus. Vastus medialis, gluteus medialis and biceps femoris played a primary role in motor control characterization when considering 4<*k<*6.

**Figure 2.**
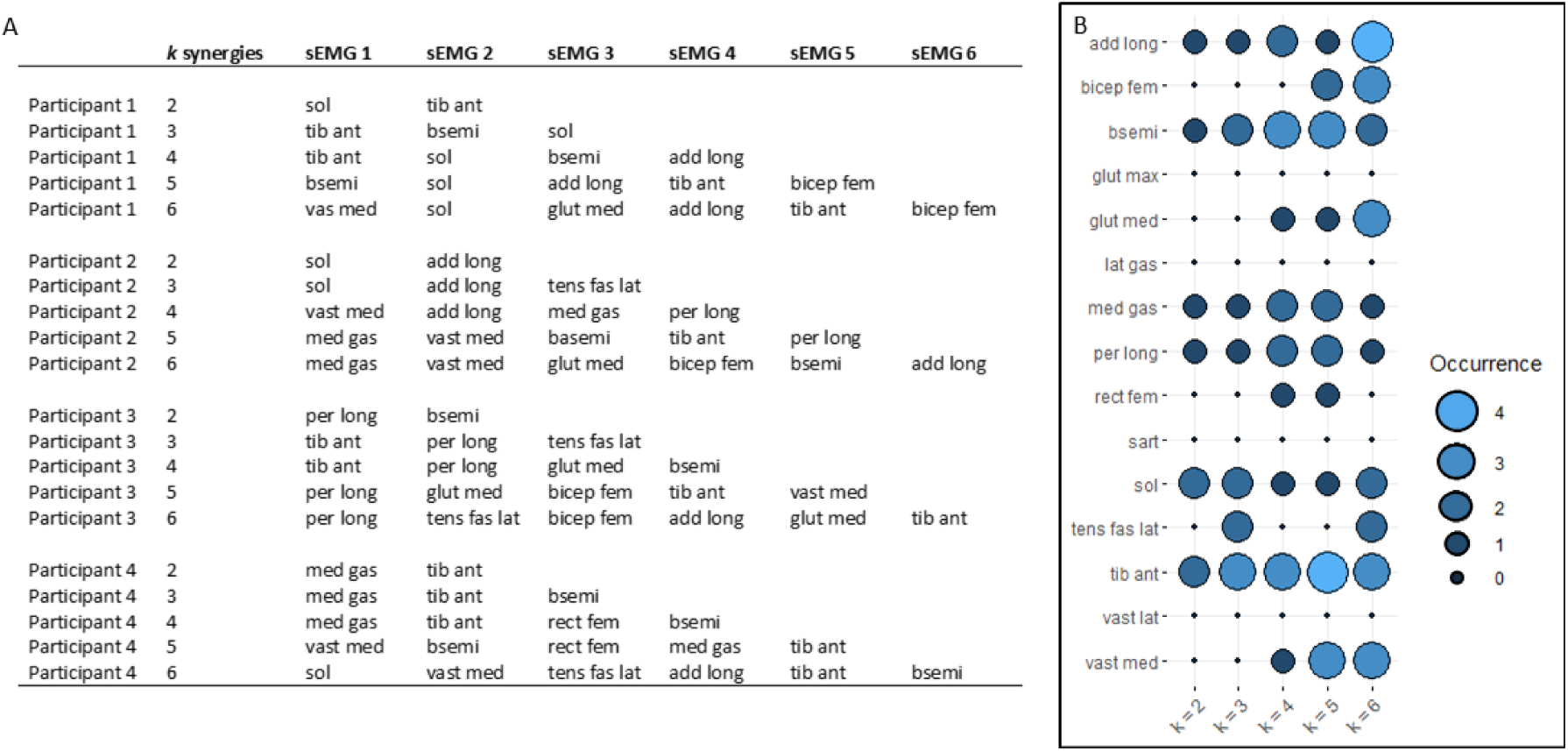
A) Table reporting which muscle excitation was chosen by the algorithm to best approximate the muscle synergy per each value of k. B) Balloon plot reporting on the occurrence of the sEMG signals used to reconstruct missing excitations, considering all the participants. The bigger the balloon, the more that specific sEMG signal has been identified to closely match to motor control patterns extracted via synergy extrapolation and thus, used in the reconstruction part.

Time series of the experimental excitations were well reproduced by the proposed method, as reported in the example (Figure 3A). VAF values on the reconstructed sEMG reported a significant difference between the investigated methods. However, the VAF value was above 95% for each subset of *k* experimental excitations used for the reconstruction (Figure 3B). Furthermore, an increase in the number of experimental excitations used for the reconstruction was associated with the highest values of VAF. The proposed method reported reasonable reconstruction accuracy, especially when using 3 to 6 experimental excitations for the prediction. Indeed, low levels of RMSD and strong to very strong levels of R^2^ were reported (Figure 3C).

**Figure 3.**
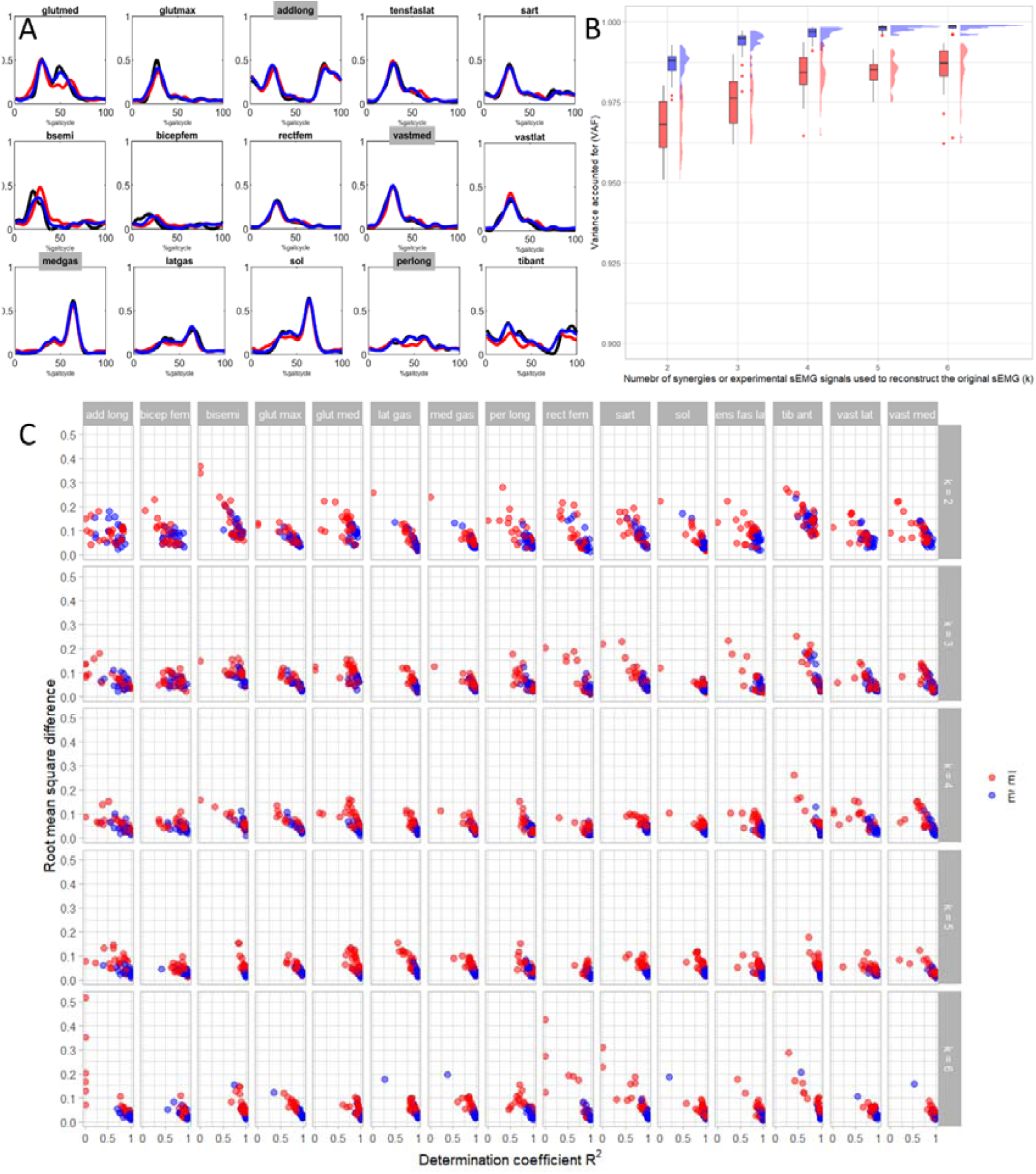
A) Excitations time series for a representative subject (mean of 6 trials). In black are reported the experimentally measured excitations, in blue the excitations reconstructed via muscle synergy analysis (using k = 4 primitives), in red the excitations reconstructed via the proposed method (using k = 4 muscles excitations as primitives, labels of the specific muscles used to predict the full set are highlighted in gray, i.e., adductor longus, vastus medialis, gastrocnemius lateralis and peroneus longus). B) Raincloud plots showing the distribution of the variance accounted for between the two methods and considering each value of k. C) Scatterplot reporting the root mean square difference in function of the determination coefficient per each reconstructed muscle excitations and each k with the two methods, in blue with the actual synergy-based methods, in red with the proposed method.

### 3.2. Neuromusculoskeletal modeling results

#### Experimental torques reconstruction accuracy

Regarding the ability of the neuromusculoskeletal models to reproduce the experimental joint moments (Figure 4), it was worth observing that acceptable performances were obtained for the knee flexion-extension moment and ankle plantar-dorsiflexion moment when using models that implied 2 or more experimental sEMG to reconstruct the whole set of used excitations. However, the models behave poorly when considering hip flexion-extension moments. This is also reflected in the metrics calculated and reported on supplementary material Table SM1.

**Figure 4.**
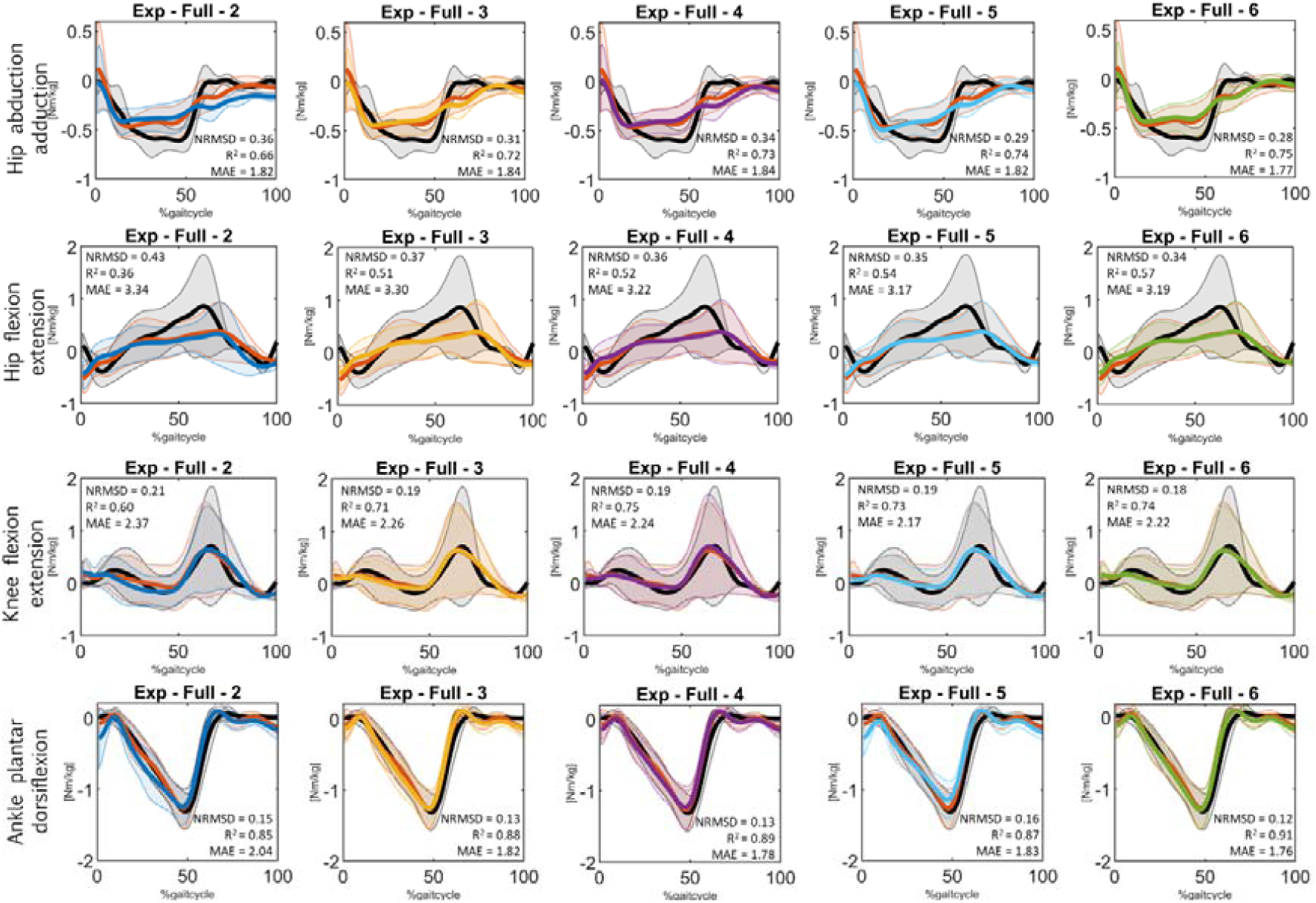
Joint torques (mean ± 1 standard deviation, solid line, and shaded area respectively). Experimental joint torques from inverse dynamics are reported in black. Simulated joint torques from the sEMG driven model with the full set of experimental sEMG are reported in red. Simulated joint torques from the sEMG driven model with 2 experimental sEMG used to simulate the full set of excitations are reported in blue. Simulated joint torques from the sEMG driven model with 3 experimental sEMG used to simulate the full set of excitations are reported in yellow. Simulated joint torques from the sEMG driven model with 4 experimental sEMG used to simulate the full set of excitations are reported in pink. Simulated joint torques from the sEMG driven model with 5 experimental sEMG used to simulate the full set of excitations are reported in cyan. Simulated joint torques from the sEMG driven model with 6 experimental sEMG used to simulate the full set of excitations are reported in green. Values of the considered metrics between experimental and simulated torques are reported in each subfigure.

#### Muscle forces prediction

Time series of the predicted muscle forces with each considered model are reported (Figure 5). Models with lower numbers of experimental sEMG used to predict the full set of muscle excitations displayed worse performances with respect to models employing more sEMG signals. Indeed, models using 2 or 3 sEMG signals to reconstruct the missing muscle excitations, reported statistically significant differences in MAE, NRMSD and R^2^ metrics with respect to models using 4, 5 or 6 sEMG signals. Muscles most affected by modeling choices were the gluteus medius, the quadriceps and the hamstrings (supplementary material, Table SM1).

**Figure 5.**
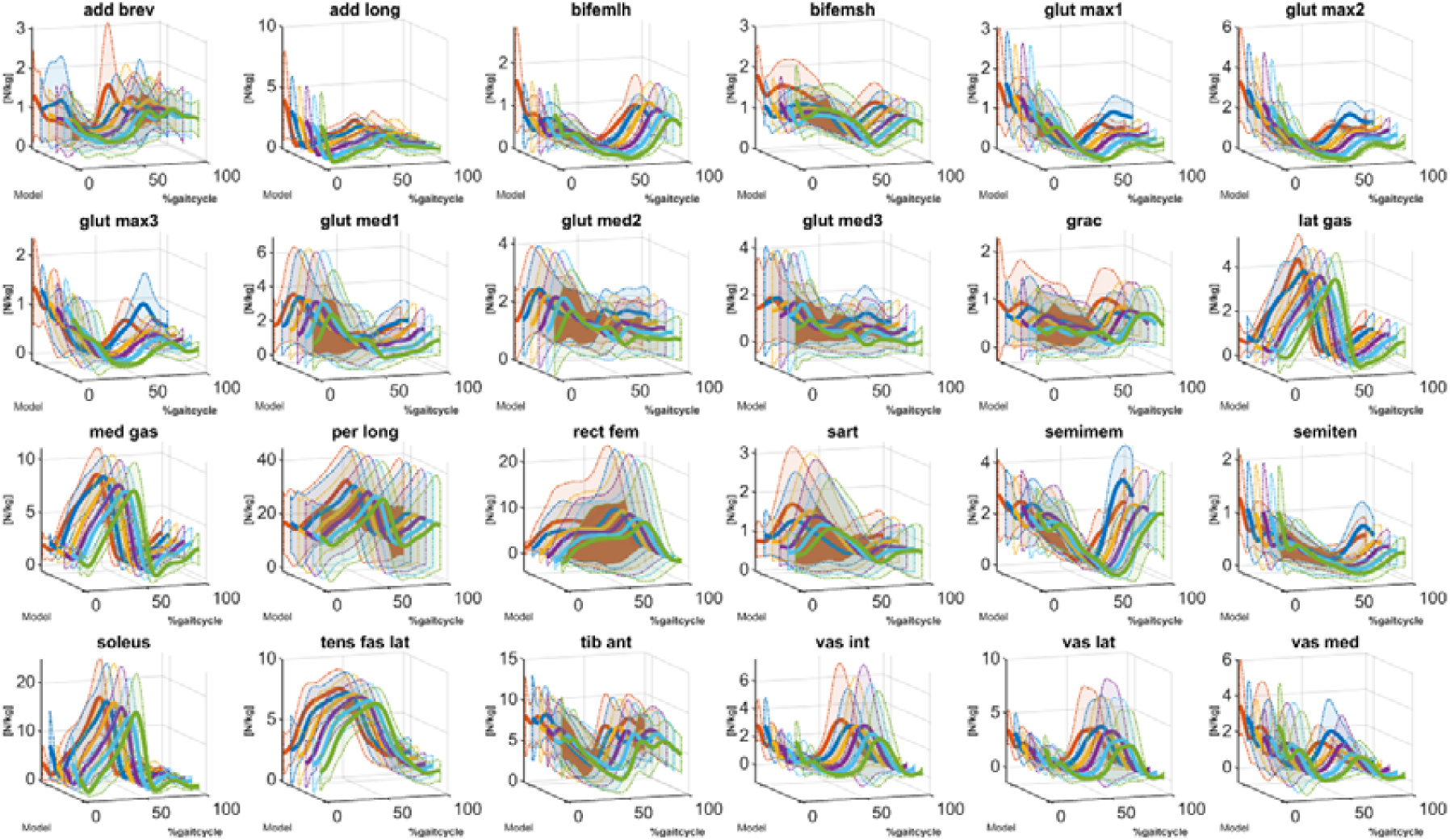
Estimated muscle forces (mean, in dotted line). Simulated muscle forces with a sEMG driven model with the full set of experimental sEMG are reported in red (gold standard model). Simulated muscle forces from a sEMG driven model with 2 experimental sEMG used to simulate the full set of excitations are reported in blue. Simulated muscle forces from a sEMG driven model with 3 experimental sEMG used to simulate the full set of excitations are reported in yellow. Simulated muscle forces from a sEMG driven model with 4 experimental sEMG used to simulate the full set of excitations are reported in purple. Simulated muscle forces from a sEMG driven model with 5 experimental sEMG used to simulate the full set of excitations are reported in cyan. Simulated muscle forces from a sEMG driven model with 6 experimental sEMG used to simulate the full set of excitations are reported in green.

## 4. DISCUSSION

The main hypothesis of the present study was that few experimental myoelectric recordings were sufficient to infer on the subject-specific lower-limb neuromuscular status. This has been tested in a novel set of trials, where the previous information has been used to reconstruct unmeasured muscle excitations (from the obtained synthetized muscle excitations) and, sEMG-driven neuromusculoskeletal models were employed to predict lower-limb joint torques and muscle forces.

The reconstruction of the 15 lower-limb muscle excitations has been obtained using a reduced number (2<*k*<6) of sEMG signals which were previously defined in the calibration part, together with a subject-specific matrix of weights and a vector of scaling factors. Results (Figure 3) suggested an acceptable prediction accuracy of the proposed method, which was able to replicate the shape and the magnitude of the experimental excitations, with a growing accuracy concomitant with the increasing of *k*. Moreover, levels of VAF obtained by the proposed method are comparable with the one obtained implying actual muscle synergies for reconstruction. Indeed, values of VAF>0.95 were reported, indicating a tolerable correctness in explaining experimental measurements by the proposed method [30]. NNMF is a factorization method based on mathematical assumption and the metric chosen to select the closest match between experimental and synergies (NRMSD) is a statistical error indicator. Despite that, the sEMGs chosen to represent motor control primitives in the synergy-based parameter calibration step, were associated with major flexor/extensor muscle contributors during gait (Figure 2A) [31], thus suggesting a physiological validity of the approach [10,11]. The chosen muscle excitations provided the necessary flexor/extensor joint torques as illustrated in Figure 4 and reported in Table SM1. Strong correlations with experimental data, together with low values of RMSD, were reported especially when considering 4 to 6 experimental sEMG signals for reconstruction purposes. Consequently, adopting a lower number of experimental sEMG could affect the reconstruction accuracy and lead to results weakly correlated with the experimental data.

Many works proposed synergy extrapolation as a method to predict missing muscle excitation from muscle synergy information derived from measured sEMG signals [12,32]. Yet, this framework was validated when just one muscle was missing (i.e., hip muscle in the specific case) and authors acknowledged that the approach should be evaluated in situations where multiple muscle recordings are missing. With our work we tried to address this issue, presenting a method able to synthetize up to 15 muscle excitations using 2 to 6 sEMG recordings. A similar attempt was done by [33] proposing a method to model the synergy function as a gaussian process. Where an approximation of their behavior was developed to allow estimation of unmeasured muscle excitations from a measured subset of sEMG. However, a major limitation of this work, as specified by the authors themselves, is the large computational time (∼31 minutes per subject) when using gaussian process regression in the model training. Adopting our proposed approach, a few minutes are enough to run both the synergy-based parameters calibration step and the neuromusculoskeletal model parameters calibration.

Moving a step further, considering the results obtained using the reconstructed muscle excitations in neuromusculoskeletal modeling framework, we found promising results in hip abduction\adduction, ankle and knee flexion\extension joint torques in terms of the extracted metrics and shapes (supplementary material Table SM1, Figure 4). Unsatisfactory results were obtained in hip flexion\extension joint torque, where an error higher than 35% was reached across every condition. This could be explained by the absence of any experimental information in major deep hip muscles such as iliopsoas and psoas [34]. Concerning muscle forces estimation, it was interesting to notice that statistically significant differences between models adopting 4 to 6 sEMG for excitation reconstruction were not found in the proposed metrics, suggesting a comparable level of accuracy among the three different models.

This study comes with several limitations that must be acknowledged. The primary one is that the method has only been tested on a small sample of non-disabled adults who exhibit no signs of disrupted motor control. Consequently, we cannot extrapolate its generalizability or applicability to individuals with neurodegenerative disorders. As a result, our statistical analysis did not account for multiple observations of the same subjects. Given the exploratory nature of our study, the sample size was insufficient to perform mixed-effects statistical models or MANOVA.

Moreover, we only assessed the participants’ preferred walking speed, without considering various walking conditions or tasks. However, the results of our preliminary analysis have provided valuable insights that support the proposed methodology. Therefore, future research should concentrate on expanding the sample size, including individuals with neurological disorders performing different tasks at varying speeds, and incorporating multiple repetitions in the statistical tests. Secondly, methodological choices had to be made to implement an automatic and consistent pipeline for data processing analysis and different choices could have led to different results (Banks et al., 2017). Indeed, muscle synergies have been extracted using NNMF methods, while different studies investigating similar topics used principal component analysis [12,32]. However, we based our choices on the work of [35], which proved that NNMF is the most appropriate method for identifying muscle synergies in walking tasks. Furthermore, *k* was constrained to vary to a limited set of numbers. However, it is widely reported in the state-of-the-art that 2 to 6 muscle synergies could be adopted to obtain reasonable information on motor control [16,26]. Additionally, the number of MTUs considered in the neuromusculoskeletal model was lower than the number of synthetized sEMG excitations used to inform the MTUs’ activation dynamics. Authors chose to use the same excitation for MTUs of the same muscle group sharing similar anatomical functions, as suggested in [36]. Finally, more research work is needed to determine how different walking conditions and methodological choices could impact on the results.

## 5. CONCLUSION

The proposed method explores the possibility to define an ecological setup for neuromusculoskeletal modeling applicable in out-of-the-lab conditions where there is the need to cope with a reduced number of sensors maintaining a reliable characterization of the individual’s neuromuscular status. Thus, in this proof-of-concept, a set of electrophysiological measurements that can be retained as surrogate of the participant’s motor control has been sought. Future developments should focus on the introduction of presented methodology in a tele-rehabilitative environment where trainings would be based on meaningful biomechanical biomarkers (*i.e., muscle forces*) that objectively report on the motor capacity of the patient, reducing possible sources of variability affecting clinical scales.

## Acknowledgements

This work was supported by MIUR (Italian Minister for Education, Universities and Research), under the initiatives of “Departments of Excellence” (Law 232/2016).

