## Supplementary table for "Muscle synergies to reduce the number of electromyography channels in neuromusculoskeletal modelling: a pilot study"

| | | Experimental recordings used to reconstruct muscle excitations | | | | |
|---|---|---|---|---|---|---|
| | | *2 sEMG* | *3 sEMG* | *4 sEMG* | *5 sEMG* | *6 sEMG* |
| *Muscle tendon unit* | *Metric* | | | | | |
| Adductor longus | *MAE* | 4.08 [0.37 – 12.61] | 3.99 [0.46 – 12.44] | 3.33 [0.23 – 9.34] | 3.59 [0.29 – 12.78] | 2.79 [0.33 – 9.72] |
| | *NRMSD* | 0.14 [0.04 – 0.67] | 0.15 [0.03 – 0.50] | 0.14 [0.01 – 0.78] | 0.14 [0.02 – 0.44] | 0.11 [0.02 – 0.23] |
| | $R^2$ | 0.87 [0.49 – 0.99] | 0.83 [0.08 – 0.99] | 0.89 [0.68 – 0.99] | 0.87 [0.54 – 0.99] | 0.89 [0.56 – 0.99] |
| Adductor brevis | *MAE* | 4.57 [0.19 – 21.68] | 3.27 [0.19 – 12.93] | 3.24 [0.17 – 12.56] | 3.97 [0.16 – 11.07] | 2.76 [0.16 – 9.21] |
| | *NRMSD* | 0.27 [0.07 – 0.49] | 0.21 [0.02 – 0.49] | 0.23 [0.06 – 0.46] | 0.22 [0.05 – 0.46] | 0.20 [0.05 – 0.45] |
| | $R^2$ | 0.78 [0.00 – 0.99] | 0.83 [0.00 – 0.99] | 0.87 [0.25 – 0.99] | 0.87 [0.53 – 0.99] | 0.78 [0.00 – 0.99] |
| Gracilis | *MAE* | 1.17 [0.19 – 2.76] | 1.13 [0.29 – 3.27] | 1.00 [0.27 – 2.37] [5] | 1.79 [0.19 – 4.97] [6] | 0.95 [0.31 – 2.22] |
| | *NRMSD* | 0.36 [0.09 – 0.87] | 0.33 [0.05 – 0.74] | 0.32 [0.04 – 0.91] | 0.34 [0.06 – 1.31] | 0.26 [0.04 – 0.62] |
| | $R^2$ | 0.76 [0.00 – 0.95] | 0.72 [0.00 – 0.99] | 0.81 [0.00 – 0.99] | 0.82 [0.00 – 0.99] | 0.73 [0.00 – 0.99] |
| Sartorius | *MAE* | 1.83 [0.07 – 3.46] | 1.55 [0.07 – 3.04] | 1.22 [0.03 – 2.99] | 1.45 [0.03 – 2.98] | 1.47 [0.03 – 2.84] |
| | *NRMSD* | 0.26 [0.08 – 0.67] | 0.21 [0.06 – 0.59] | 0.15 [0.04 – 0.42] | 0.21 [0.07 – 1.19] | 0.17 [0.04 – 0.37] |
| | $R^2$ | 0.87 [0.63 – 0.99] | 0.87 [0.00 – 0.99] | 0.93 [0.77 – 0.99] | 0.94 [0.77 – 0.99] | 0.94 [0.69 – 0.99] |
| Tensor fascia latae | *MAE* | 2.11 [0.67 – 4.83] | 1.47 [0.12 – 4.20] | 1.42 [0.25 – 3.62] | 1.27 [0.20 – 3.57] | 1.18 [0.17 – 2.79] |
| | *NRMSD* | 0.07 [0.02 – 0.22] | 0.05 [0.00 – 0.23] | 0.04 [0.00 – 0.19] | 0.03 [0.01 – 0.12] | 0.04 [0.00 – 0.11] |
| | $R^2$ | 0.99 [0.92 – 0.99] | 0.99 [0.91 – 1.00] | 0.99 [0.96 – 1.00] | 0.99 [0.96 – 1.00] | 0.99 [0.98 – 1.00] |
| Gluteus medius anterior | *MAE* | 4.01 [1.15 – 7.56] [6] | 3.45 [1.09 – 6.63] [6] | 3.09 [0.72 – 6.59] | 3.01 [0.73 – 7.26] | 1.60 [0.57 – 3.54] |
| | *NRMSD* | 0.18 [0.08 – 0.29] [5, 6] | 0.15 [0.04 – 0.34] [6] | 0.14 [0.03 – 0.31] | 0.10 [0.02 – 0.20] | 0.08 [0.02 – 0.31] |
| | $R^2$ | 0.89 [0.68 – 0.98] [5, 6] | 0.91 [0.68 – 0.99] | 0.92 [0.75 – 0.99] | 0.95 [0.87 – 0.99] | 0.97 [0.82 – 0.99] |
| Gluteus medius posterior | *MAE* | 3.07 [0.82 – 7.55] [6] | 2.62 [0.81 – 5.98] [6] | 2.38 [0.70 – 5.78] | 2.23 [0.64 – 5.58] | 1.23 [0.48 – 2.80] |
| | *NRMSD* | 0.29 [0.12 – 0.49] [4, 5, 6] | 0.22 [0.08 – 0.53] [5, 6] | 0.19 [0.05 – 0.39] | 0.14 [0.04 – 0.32] | 0.11 [0.03 – 0.29] |
| | $R^2$ | 0.86 [0.63 – 0.98] [5, 6] | 0.90 [0.54 – 0.99] [6] | 0.92 [0.75 – 0.99] | 0.95 [0.78 – 0.99] | 0.98 [0.86 – 0.99] |
| Gluteus medius middle | *MAE* | 4.21 [0.58 – 8.87] [6] | 3.39 [0.55 – 7.02] [6] | 2.92 [0.61 – 7.00] | 2.84 [0.48 – 6.29] | 1.56 [0.43 – 3.28] |
| | *NRMSD* | 0.29 [0.09 – 0.63] [4, 5, 6] | 0.55 [0.08 – 0.60] [6] | 0.18 [0.04 – 0.49] | 0.15 [0.04 – 0.39] | 0.12 [0.04 – 0.27] |
| | $R^2$ | 0.71 [0.00 – 0.92] [3, 4, 5, 6] | 7.02 [0.61 – 0.97] | 0.86 [0.52 – 0.99] | 0.88 [0.58 – 0.99] | 0.95 [0.82 – 0.99] |
| Gluteus maximus superior | *MAE* | 1.91 [0.33 – 3.59] [4, 5] | 1.49 [0.32 – 2.60] | 1.29 [0.29 – 2.69] | 1.26 [0.22 – 2.66] | 1.34 [0.25 – 2.88] |
| | *NRMSD* | 0.20 [0.07 – 0.46] | 0.18 [0.03 – 0.45] | 0.20 [0.03 – 0.67] | 0.15 [0.03 – 0.45] | 0.18 [0.03 – 0.46] |
| | $R^2$ | 0.79 [0.31 – 0.96] [5] | 0.87 [0.65 – 0.98] | 0.86 [0.52 – 0.99] | 0.89 [0.68 – 0.98] | 0.88 [0.65 – 0.99] |
| Gluteus maximus medialis | *MAE* | 3.27 [0.80 – 6.97] [4, 5, 6] | 2.28 [0.23 – 4.50] | 2.06 [0.23 – 3.84] | 1.97 [0.24 – 4.39] | 1.96 [0.34 – 5.08] |
| | *NRMSD* | 0.18 [0.06 – 0.40] | 0.15 [0.03 – 0.37] | 0.18 [0.02 – 0.56] | 0.13 [0.04 – 0.35] | 0.15 [0.04 – 0.36] |
| | $R^2$ | 0.79 [0.32 – 0.95] [5, 6] | 0.88 [0.53 – 0.99] | 0.87 [0.58 – 0.99] | 0.91 [0.76 – 0.99] | 0.91 [0.72 – 0.99] |
| Gluteus maximus inferior | *MAE* | 2.69 [0.59 – 7.32] | 2.09 [0.42 – 5.37] | 1.84 [0.36 – 4.46] | 1.81 [0.32 – 5.09] | 1.78 [0.33 – 4.77] |
| | *NRMSD* | 0.22 [0.07 – 0.33] | 0.18 [0.05 – 0.31] | 0.18 [0.03 – 0.45] | 0.15 [0.03 – 0.26] | 0.18 [0.03 – 0.45] |
| | $R^2$ | 0.68 [0.00 – 0.95] [3, 4, 5, 6] | 0.84 [0.56 – 0.96] | 0.84 [0.55 – 0.99] | 0.87 [0.61 – 0.98] | 0.87 [0.63 – 0.98] |
| Biceps femoris long head | *MAE* | 2.36 [0.36 – 4.88] [6] | 2.05 [0.17 – 5.62] | 1.89 [0.24 – 5.55] | 1.35 [0.20 – 4.23] | 1.06 [0.29 – 2.38] |
| | *NRMSD* | 0.29 [0.07 – 0.52] [6] | 0.26 [0.05 – 0.53] [6] | 0.24 [0.06 – 0.56] [6] | 0.19 [0.03 – 0.61] | 0.14 [0.06 – 0.28] |
| | $R^2$ | 0.72 [0.29 – 0.95] [6] | 0.74 [0.16 – 0.98] [6] | 0.77 [0.11 – 0.98] | 0.86 [0.44 – 0.99] | 0.91 [0.75 – 0.99] |

| | | | | | | |
|---|---|---|---|---|---|---|
| Biceps femoris short head | *MAE* | 1.14 [0.42 – 1.97] [5, 6] | 1.02 [0.26 – 1.86] [6] | 0.95 [0.22 – 2.12] | 0.75 [0.11 – 1.68] | 0.61 [0.22 – 1.43] |
| | *NRMSD* | 0.37 [0.06 – 1.25] [6] | 0.31 [0.06 – 0.95] | 0.31 [0.05 – 0.98] | 0.27 [0.01 – 0.79] | 0.16 [0.03 – 0.48] |
| | $R^2$ | 0.89 [0.69 – 0.99] [6] | 0.89 [0.68 – 0.99] | 0.89 [0.66 – 0.99] | 0.93 [0.67 – 0.99] | 0.95 [0.80 – 0.99] |
| Semimembranosus | *MAE* | 5.14 [1.29 – 12.88] [5] | 4.47 [0.32 – 14.05] | 3.70 [0.34 – 14.29] | 2.04 [0.41 – 5.91] | 2.68 [0.57 – 7.78] |
| | *NRMSD* | 0.19 [0.07 – 0.29] [4, 5] | 0.16 [0.02 – 0.49] | 0.11 [0.02 – 0.43] | 0.10 [0.03 – 0.41] | 0.15 [0.05 – 0.39] |
| | $R^2$ | 0.77 [0.37 – 0.98] [3, 4, 5, 6] | 0.88 [0.34 – 0.99] | 0.90 [0.21 – 0.99] | 0.95 [0.77 – 0.99] | 0.89 [0.70 – 0.99] |
| Semitendinosus | *MAE* | 1.16 [0.27 – 2.54] [3, 4, 5, 6] | 0.65 [0.09 – 1.65] | 0.58 [0.02 – 1.77] | 0.41 [0.07 – 0.89] | 0.66 [0.09 – 1.71] |
| | *NRMSD* | 0.17 [0.06 – 0.38] [5, 6] | 0.15 [0.01 – 0.43] | 0.09 [0.01 – 0.38] | 0.07 [0.01 – 0.27] | 0.09 [0.02 – 0.23] |
| | $R^2$ | 0.84 [0.66 – 0.98] [5] | 0.87 [0.35 – 0.99] | 0.88 [0.30 – 0.99] | 0.96 [0.78 – 0.99] | 0.93 [0.68 – 0.99] |
| Rectus femoris | *MAE* | 7.07 [0.76 – 23.24] [3, 4, 5] | 3.79 [0.99 – 9.02] | 3.29 [0.98 – 9.77] | 3.01 [1.29 – 11.98] | 4.41 [0.96 – 11.58] |
| | *NRMSD* | 0.20 [0.05 – 0.64] [4, 5, 6] | 0.13 [0.01 – 0.39] | 0.09 [0.02 – 0.20] | 0.11 [0.02 – 0.18] | 0.11 [0.01 – 0.29] |
| | $R^2$ | 0.89 [0.61 – 0.99] | 0.91 [0.12 – 0.99] | 0.97 [0.79 – 0.99] | 0.97 [0.91 – 0.99] | 0.95 [0.59 – 0.99] |
| Vastus medialis | *MAE* | 5.07 [1.21 – 16.35] | 3.51 [0.64 – 9.71] | 2.94 [1.10 – 8.06] | 1.98 [0.49 – 4.89] | 1.79 [0.36 – 5.63] |
| | *NRMSD* | 0.19 [0.10 – 0.35] | 0.16 [0.04 – 0.35] | 0.16 [0.04 – 0.35] | 0.13 [0.02 – 0.33] | 0.07 [0.01 – 0.22] |
| | $R^2$ | 0.76 [0.47 – 0.93] | 0.86 [0.67 – 0.99] | 0.84 [0.67 – 0.99] | 0.89 [0.44 – 0 .99] | 0.97 [0.88 – 0.99] |
| Vastus lateralis | *MAE* | 5.22 [1.58 – 15.79] [3, 5, 6] | 3.24 [1.15 – 7.81] | 4.45 [1.18 – 12.34] | 3.12 [0.77 – 7.27] | 2.89 [1.17 – 10.09] |
| | *NRMSD* | 0.17 [0.05 – 0.31] [6] | 0.13 [0.03 – 0.26] | 0.17 [0.03 – 0.39] [6] | 0.12 [0.03 – 0.33] | 0.08 [0.04 – 0.16] |
| | $R^2$ | 0.73 [0.00 – 0.98] [3, 5, 6] | 0.88 [0.66 – 0.99] | 0.79 [0.23 – 0.99] [6] | 0.89 [0.64 – 0.99] | 0.93 [0.75 – 0.99] |
| Vastus intermedius | *MAE* | 4.08 [0.87 – 14.28] [3, 5, 6] | 2.26 [0.41 – 4.71] | 2.58 [0.66 – 4.82] | 1.80 [0.17 – 5.57] | 1.63 [0.41 – 7.46] |
| | *NRMSD* | 0.17 [0.06 – 0.37] [6] | 0.13 [0.02 – 0.31] | 0.16 [0.02 – 0.43] [6] | 0.11 [0.01 – 0.32] | 0.06 [0.01 – 0.12] |
| | $R^2$ | 0.79 [0.25 – 0.98] [3, 5, 6] | 0.90 [0.73 – 0.99] | 0.85 [0.44 – 0.99] [6] | 0.91 [0.67 – 0.99] | 0.97 [0.88 – 0.99] |
| Gastrocnemius lateralis | *MAE* | 3.25 [0.76 – 11.00] | 2.58 [0.69 – 5.87] | 2.62 [0.96 – 5.35] | 3.01 [0.86 – 6.59] | 2.57 [0.94 – 5.28] |
| | *NRMSD* | 0.11 [0.01 – 0.29] | 0.08 [0.01 – 0.23] | 0.09 [0.01 – 0.22] | 0.13 [0.01 – 0.39] | 0.08 [0.01 – 0.20] |
| | $R^2$ | 0.90 [0.49 – 0.99] | 0.95 [0.74 – 0.99] | 0.95 [0.73 – 0.99] | 0.91 [0.66 – 0.99] | 0.94 [0.70 – 0.99] |
| Gastrocnemius medialis | *MAE* | 4.59 [1.62 – 19.61] | 3.60 [1.68 – 10.46] | 3.23 [1.13 – 9.37] | 4.21 [0.78 – 7.55] | 3.73 [1.09 – 7.12] |
| | *NRMSD* | 0.09 [0.02 – 0.31] | 0.07 [0.01 – 0.27] | 0.06 [0.01 – 0.25] | 0.09 [0.01 – 0.22] | 0.07 [0.01 – 0.19] |
| | $R^2$ | 0.89 [0.00 – 0.99] | 0.92 [0.00 – 0.99] | 0.93 [0.09 – 0.99] | 0.94 [0.45 – 0.99] | 0.95 [0.59 – 0.99] |
| Peroneus longus | *MAE* | 9.78 [1.95 – 22.39] [3, 4, 5, 6] | 6.20 [1.99 – 10.24] | 5.03 [0.81 – 10.47] | 5.99 [1.83 – 12.03] | 6.26 [2.06 – 12.61] |
| | *NRMSD* | 0.07 [0.01 – 0.16] [4] | 0.04 [0.02 – 0.16] | 0.04 [0.01 – 0.19] | 0.04 [0.01 – 0.19] | 0.05 [0.02 – 0.21] |
| | $R^2$ | 0.98 [0.92 – 0.99] | 0.99 [0.93 – 0.99] | 0.99 [0.89 – 1.00] | 0.99 [0.89 – 0.99] | 0.99 [0.89 – 0.99] |
| Soleus | *MAE* | 13.00 [2.02 – 41.07] | 8.68 [2.17 – 29.42] | 11.04 [3.89 – 28.79] | 12.56 [7.19 – 23.98] | 9.78 [2.04 – 26.13] |
| | *NRMSD* | 0.08 [0.01 – 0.22] | 0.06 [0.02 – 0.16] [5] | 0.11 [0.03 – 0.44] | 0.14 [0.04 – 0.41] | 0.08 [0.01 – 0.24] |
| | $R^2$ | 0.92 [0.55 – 0.99] | 0.97 [0.75 – 0.99] | 0.93 [0.77 – 0.99] | 0.92 [0.78 – 0.99] | 0.96 [0.86 – 0.99] |
| Tibialis anterior | *MAE* | 10.37 [4.88 – 25.72] [3, 4, 5, 6] | 8.91 [1.19 – 25.07] | 7.63 [0.99 – 32.28] | 6.69 [1.94 – 17.86] | 7.97 [2.19 – 27.85] |
| | *NRMSD* | 0.18 [0.04 – 0.44] | 0.13 [0.02 – 0.42] | 0.11 [0.02 – 0.45] | 0.12 [0.04 – 0.35] | 0.13 [0.02 – 0.49] |
| | $R^2$ | 0.89 [0.47 – 0.98] | 0.97 [0.89 – 0.99] | 0.98 [0.88 – 0.99] | 0.98 [0.94 – 0.99] | 0.97 [0.85 – 0.99] |

| | | | | | | |
|---|---|---|---|---|---|---|
| | | | *Experimental measures used to reconstruct the excitations* | | | |
| *Joint torques* | | *2 sEMG* | *3 sEMG* | *4 sEMG* | *5 sEMG* | *6 sEMG* |

| | | | | | | |
|---|---|---|---|---|---|---|
| Hip Abduction – Adduction | *MAE* | 1.82 [1.29 – 2.53] | 1.84 [1.25 – 2.59] | 1.84 [1.25 – 2.69] | 1.82 [1.28 – 2.66] | 1.78 [1.21 – 2.65] |
| | *NRMSD* | 0.36 [0.17 – 0.68] | 0.31 [0.13 – 0.69] | 0.34 [0.14 – 0.78] | 0.29 [0.13 – 0.58] | 0.28 [0.14 – 0.53] |
| | $R^2$ | 0.66 [0.20 – 0.86] | 0.73 [0.22 – 0.88] | 0.73 [0.45 – 0.87] | 0.74 [0.47 – 0.89] | 0.75 [0.42 – 0.89] |
| Hip Flexion – Extension | *MAE* | 3.35 [1.84 – 7.00] | 3.30 [1.86 – 6.99] | 3.23 [1.68 – 6.80] | 3.17 [1.84 – 6.45] | 3.19 [1-81 – 6.63] |
| | *NRMSD* | 0.43 [0.18 – 0.77] | 0.37 [0.20 – 0.56] | 0.36 [0.14 – 0.60] | 0.35 [0.16 – 0.47] | 0.35 [0.22 – 0.47] |
| | $R^2$ | 0.36 [0.00 – 0.65] $^{3, 4, 5, 6}$ | 0.51 [0.13 – 0.71] | 0.52 [0.19 – 0.79] | 0.54 [0.18 – 0.73] | 0.57 [0.39 – 0.68] |
| Knee Flexion – Extension | *MAE* | 2.37 [1.02 – 5.83] | 2.27 [0.93 – 5.67] | 2.24 [0.89 – 5.72] | 2.17 [0.82 – 5.34] | 2.22 [0.89 – 5.40] |
| | *NRMSD* | 0.21 [0.14 – 0.30] | 0.19 [0.11 – 0.32] | 0.19 [0.09 – 0.30] | 0.19 [0.10 – 0.28] | 0.18 [0.10 – 0.27] |
| | $R^2$ | 0.60 [0.00 – 0.87] | 0.71 [0.32 – 0.89] | 0.75 [0.52 – 0.93] | 0.73 [0.34 – 0.92] | 0.73 [0.44 – 0.91] |
| Ankle Plantar – Dorsiflexion | *MAE* | 2.04 [1.31 – 3.39] | 1.82 [1.91 – 3.18] | 1.78 [1.19 – 3.08] | 1.83 [1.11 – 2.84] | 1.76 [1.04 – 3.16] |
| | *NRMSD* | 0.15 [0.07 – 0.23] | 0.13 [0.06 – 0.20] | 0.13 [0.06 – 0.23] | 0.16 [0.04 – 0.32] | 0.12 [0.05 – 0.22] |
| | $R^2$ | 0.85 [0.46 – 0.96] | 0.89 [0.68 – 0.98] | 0.89 [0.67 – 0.98] | 0.87 [0.72 – 0.99] | 0.91 [0.67 – 0.98] |

*Table SM1.* Values of the considered metrics against the experimental and the simulated measures. The mean, the lowest and the highest (in square brackets) value of mean absolute error (MAE), normalized root mean square difference (NRMSD), and determination coefficient ($R^2$) are reported for muscle forces and joint torques obtained by each model. Statistically significant differences ($p < 0.05$) between models are reported with a number as apex. The number indicates the model that is statistically different form the considered one.
